# AGO CLIP reveals an activated network for acute regulation of brain glutamate homeostasis after ischemic stroke

**DOI:** 10.1101/245928

**Authors:** Mariko Kobayashi, Corinne Benakis, Corey Anderson, Michael J. Moore, Carrie Poon, Ken Uekawa, Jonathan P. Dyke, John J. Fak, Aldo Mele, Christopher Y. Park, Ping Zhou, Josef Anrather, Costantino Iadecola, Robert B. Darnell

## Abstract

Post-transcriptional regulation by miRNAs is essential for complex molecular responses to physiological insult and disease. Although many disease-associated miRNAs are known, their global targets and culminating network effects on pathophysiology remain poorly understood. We applied AGO CLIP to systematically elucidate altered miRNA-target interactions in brain following ischemia/reperfusion (I/R) injury. Among 1,190 identified, most prominent was the cumulative loss of target regulation by miR-29 family members. Integration of translational and time-course RNA profiles revealed a dynamic mode of miR-29 target de-regulation, led by acute translational activation and later increase in RNA levels, allowing rapid proteomic changes to take effect. These functional regulatory events rely on canonical and non-canonical miR-29 binding and engage glutamate reuptake signals to control local glutamate levels. These results uncover a miRNA target network that acts acutely to maintain brain homeostasis after ischemic stroke.

## INTRODUCTION

Divergent post-transcriptional gene regulation in the brain is indispensable to achieve its highly specialized and diverse functions. An aspect of this control is mediated by miRNAs, small non-coding RNAs that fine-tune gene expression by base pairing to 3’UTRs for target mRNA repression (Baek et al., 2008; Bartel, 2009). A single miRNA can have multiple targets, providing coordinate governance over a network of genes. Their dynamic regulatory capacity, including precise temporal responses to external cues (Ebert and Sharp, 2012) and spatial-specific targeting of localized mRNAs (Leung and Sharp, 2006) provide an elegant molecular solution to meet the brain’s unique physiological demands. Ablation of the miRNA processing machinery, *Dicer,* leads to neuronal loss, memory/behavior deficits, neurodegenerative phenotypes akin to Parkinson’s Disease (PD) and even enhancement of poly-glutamine toxicity (Bilen et al., 2006; Kim et al., 2007). Although substantial research has determined a requirement of miRNAs for proper function, the identities, targets, and collective roles of their regulatory network in disease remain largely unknown.

The development of high-throughput approaches to capture and identify cellular RNAs has catapulted the search for miRNA regulatory networks. These unbiased, highly sensitive and quantitative technologies have been especially useful for identifying miRNAs with divergent expression patterns in dynamic physiologic contexts and in disease. However, many studies are limited by their inability to define precise gene regulatory networks impacted by altered miRNA expression. HITS-CLIP (<u>h</u>igh-throughput <u>s</u>equencing of RNAs isolated by <u>c</u>ross<u>l</u>inking <u>i</u>mmuno<u>p</u>recipitation) combines rigorous biochemical methods with hi-throughput sequencing technology to identify bona fide RNA:protein interactions (Licatalosi et al., 2008). When applied with RNA-induced silencing complex (RISC) factor Argonaute (AGO), HITS-CLIP allows for empirical mapping of miRNA targeting events through isolation of endogenous AGO:miRNA:mRNA (ternary) complexes (Chi et al., 2009, Moore et al., 2014). Application of AGO HITS-CLIP, or AGO CLIP, was instrumental for discovery of novel miRNA actions in unique biological contexts, including coordinate control of viral latency, viral sequestration of host miRNAs, viral genome propagation by 3’UTR targeting (Riley et al., 2012; Luna et al., 2015; Scheel et al., 2016), and cell invasion networks in cancer (Bracken et al., 2014). Subsequent modifications of CLIP (ie. CLEAR-CLIP) utilize covalent ligation of RNAs within AGO ternary complexes to garner miRNA:mRNA chimeras for the precise identification and analysis of *in vivo* miRNA targeting events (Moore et al., 2015, Helwak et al., 2013). These methods provide powerful tools to facilitate genome-wide discovery of miRNA function in disease with potential to uncover new targets for therapy.

Stroke is a major cause of death and disability worldwide, with limited therapeutic options (Moskowitz et al, 2010). The molecular pathology of ischemic stroke is poorly understood, particularly the early gene expression changes and regulatory events that mediate neuro-protection and ischemic cell death (Moskowitz et al., 2010, Iadecola and Anrather, 2011). In the present study we employed AGO CLIP for transcriptome-wide mapping of altered miRNA targeting events in an I/R injury model of stroke. Differential profiling of AGO bound RNA distinguished over one thousand miRNA:mRNA events changing in stroke. Detailing the global molecular events lead by a single family of robustly down-regulated miRNAs, miR-29s, unveiled a dynamic and coordinate cascade of target activation, impacting mRNA translation at an early time-point, followed by a global increase in steady-state RNA levels. Unexpectedly, many of the functional miRNA binding events relied on non-canonical seed pairing events, underscoring the importance of empirically identified targets over informatic predictions. We extend these findings to provide new insights to the network of stroke-miRNA targets mediating glutamate homeostasis and to miRNA regulatory site polymorphisms as potential determinants of disease outcome, providing novel insights and strategy toward understanding human stroke disease.

## RESULTS

### Identification of AGO:RNA interactions in stroke response

To model human ischemic stroke, we used the mouse middle cerebral artery occlusion (MCAo) model (Figure 1A). This procedure relies on the transient occlusion of the middle cerebral artery (MCA), the artery most often occluded in stroke patients, and further model the molecular events of reperfusion that follow (Casals et al., 2011). For CLIP, we chose an early time-point of 3hrs (post I/R injury) in order to distinguish direct molecular consequences of stroke injury from indirect events involving secondary phenomena such as major cell death and infiltration of blood-borne inflammatory cells (Iadecola and Anrather, 2011).

**Figure 1.**
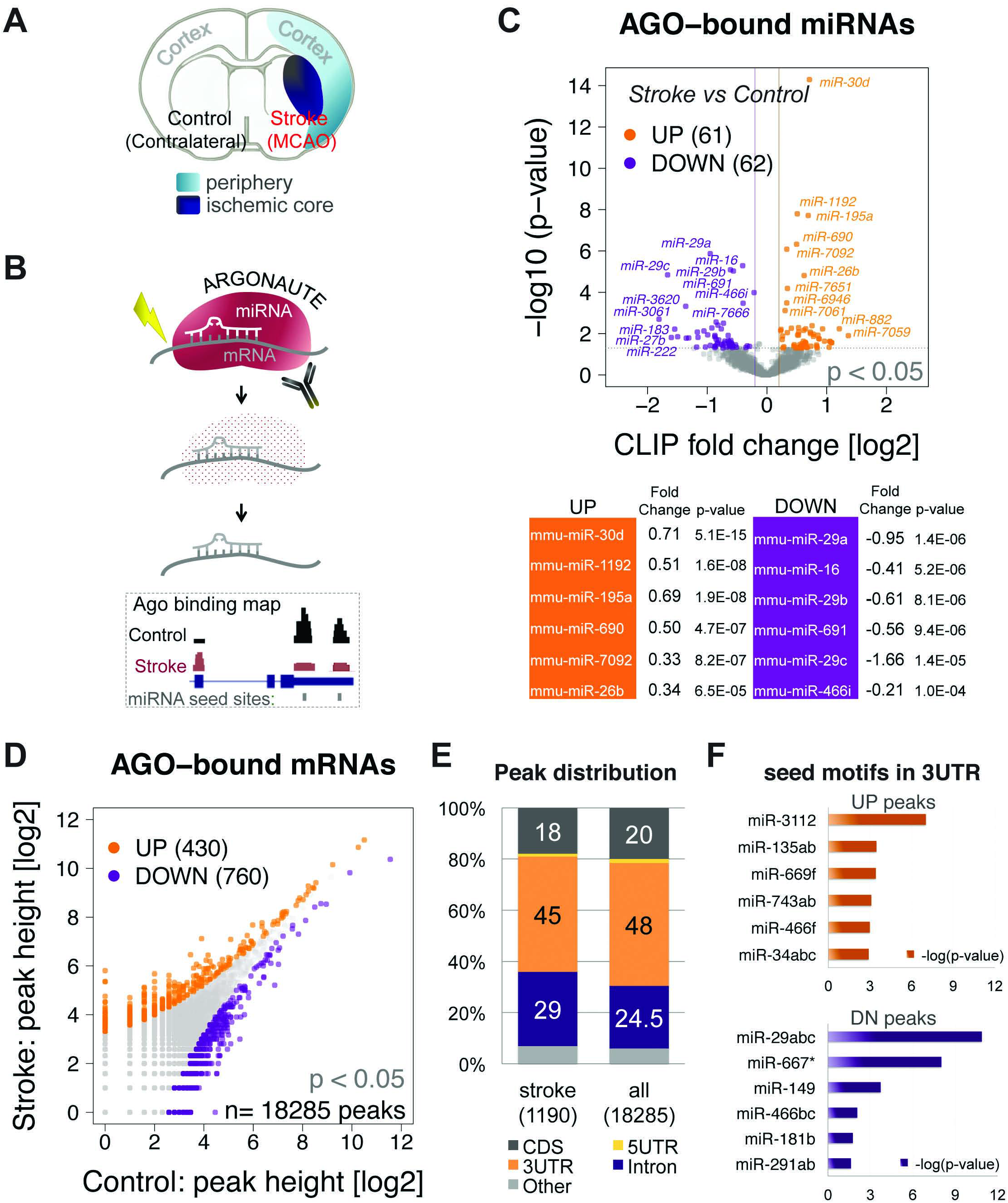
Identification of stroke-associated AGO:RNA interactions using CLIP. (A) Mouse brain coronal slice illustrating MCAo model depicting contralateral (Control) and MCAo (Stroke) hemispheres. Cortical ‘periphery’ (light blue) is distinguished from ‘ischemic core’ (dark blue). (B) Stroke AGO CLIP. 1) IP purification of UV crosslinked AGO:RNA complexes. 2) proteinase digestion of AGO 3) purification of bound RNA and 4) subsequent library generation and hi-throughput sequencing. 5) Difference analysis is performed on mapped CLIP sequencing reads. (C) Volcano plot of AGO-bound miRNAs increasing (orange) or decreasing (purple) in stroke condition. The two vertical lines indicate log2 fold change cut-off (>|0.2|) the horizontal dotted line is p-value cut-off (p<0.05). Table lists top 6 miRNAs (UP and DOWN) ranked by p-values. (D) Scatter plot of 18,285 AGO-bound mRNA peaks found in either control or stroke. Increased (orange) and decreased (purple) stroke peaks (p<0.05). (E) Distribution of peaks among annotated genomic regions for stroke peaks and for ‘all’ robust peaks. (F) Motif enrichment of 3’UTR stroke peaks, UP (top) or DN (bottom) in stroke. All stroke samples collected after 3hr I/R. See also Figure S1.

For differential analysis of AGO-associated RNA, we compared data from ‘Stroke’ hemisphere, containing the ischemic territory, to the contralateral ‘Control’ hemisphere (Figure 1A). Sham surgery animals were also assessed to rule out general affects caused by surgical anesthesia (Figure S1A). We focused our biochemical analysis on cortical tissues surrounding the center of the ischemic territory (ischemic core), the “ischemic periphery,” a region at risk for infarction targeted by intrinsic neuro-protective mechanisms (Figure 1A, Moskowitz et al., 2010). The AGO CLIP protocol on these samples was executed as previously described (Chi et al., 2009, Moore et al., 2014, Figure 1B), using stringent methods to isolate UV-crosslinked AGO:RNA complexes by immunoprecipitation and gel purification. Autoradiograms of crosslinked RNAs revealed no difference in the amount of purified complexes in the stroke samples compared to controls (Figure S1B). The isolated AGO-bound RNAs were sequenced by high-throughput sequencing and mapped to the mouse reference genome for AGO:mRNA binding peak definition, using a previously reported peak calling strategy (Chi et al., 2009, Moore et al., 2014), and to the miRBase reference of miRNAs for AGO-bound miRNA assessment (Kozomara and Griffiths-Jones, 2010).

Quantitative analysis of binding peaks, using peak height measurements (sequence read per peak) was performed to reveal stroke-specific AGO binding patterns (Figure 1B). CLIP reads across 4-5 biological replicates had high correlations (average Pearson correlation coefficients of ~0.7 to 0.9 for miRNAs, and of ~0.6 for mRNAs (Figure S1C-D). Analysis of binding events reproducible in at least 3 replicates for a single condition (biological complexity, BC3 or greater) (Moore et al., 2014) identified 123 miRNAs with significantly altered AGO association in stroke (p<0.05) of which 61 showed an increase while 62 were decreased (Figure 1C), while those identified in Sham condition showed minimal resemblance (Figure S1E). miRNAs with altered AGO binding included many that have been previously implicated in stroke, such as miR-30d, miR-26b, miR-16, miR-222, and miR-183. The most notable changes were observed among the three members of the miR-29 family, miR-29a, -29b, and -29c (Figure 1C).

For the investigation of AGO-bound mRNAs, CLIP peaks were assigned a binding region of 80 nt widths relative to peak midpoints, slightly wider than previously defined peak width (Chi et al., 2009). Only peaks containing reads from 3 out of 5 biological replicates were considered (BC≥3). Using this approach, we identified 18,285 reproducible AGO peaks, out of which 1,190 were differentially bound by AGO in stroke condition (determined by binomial testing and applying p-value cut off, p<0.05). Among these ‘stroke peaks’, 430 showed increased binding while 760 showed a significant decrease (Figure 1D). Although canonical miRNA targeting is at the 3’UTRs of mRNAs, previous CLIP studies have shown miRNA guided AGO targeting also occurs within coding sequences, 5’UTRs and in introns (Chi et al., 2009, Moore et al., 2014). To assess whether the transcriptome-wide sites of AGO targeting is altered upon I/R injury, we determined the distribution of 1,190 stroke peaks among annotated transcript regions. As expected, the majority (45%) of stroke peaks reside within annotated 3’UTR regions, while a smaller portion found in introns (29%) and coding regions (18%). We saw no difference in the genomic distribution of stroke peaks with respect to ‘all’ AGO binding events (Figure 1E). The proportion of 3’UTR peaks was higher (49%) when indeterminate binding events, also identified in ‘Sham’ condition, were removed from analysis (Figure S1F-G). Finally, we performed 6-nucleotide motif enrichment analysis of stroke peaks, among up- and down-regulated peaks within 3’UTRs, independently (Figure 1F). Strikingly, 6mer seed motif engaging miR-29 binding was the top enriched motif among down-regulated stroke peaks. Global AGO regulatory site mapping of miRNA and mRNA collectively suggest a role for miR-29 dysregulation in stroke brain.

### miR-29 family of miRNAs are down-regulated in stroke

CLIP data revealed I/R injury dependent, robust down-regulation by miR-29 family members and a synonymous loss of AGO regulation at miR-29 seed containing 3’UTRs (Fig1C, 1F). To further address this we used an orthogonal approach, measuring mature miRNA levels using small RNA sequencing. Although changes in AGO-bound miRNAs and steady-state levels were poorly correlated (Pearson correlation coefficient = −0.1), we saw a corroborative decrease for all miR-29 members in both datasets (Figure 2A, S2B). This was further confirmed by qPCR quantification of each miR-29 family member (Figure 2B). Loss of miR-29 was not detected in the sham surgery condition (Figure 2B, S2A), ruling out confounding effects of anesthesia and surgical manipulation. The data suggest that loss of AGO guided miR-29 regulation is likely due to a decrease in steady-state RNA levels.

**Figure 2.**
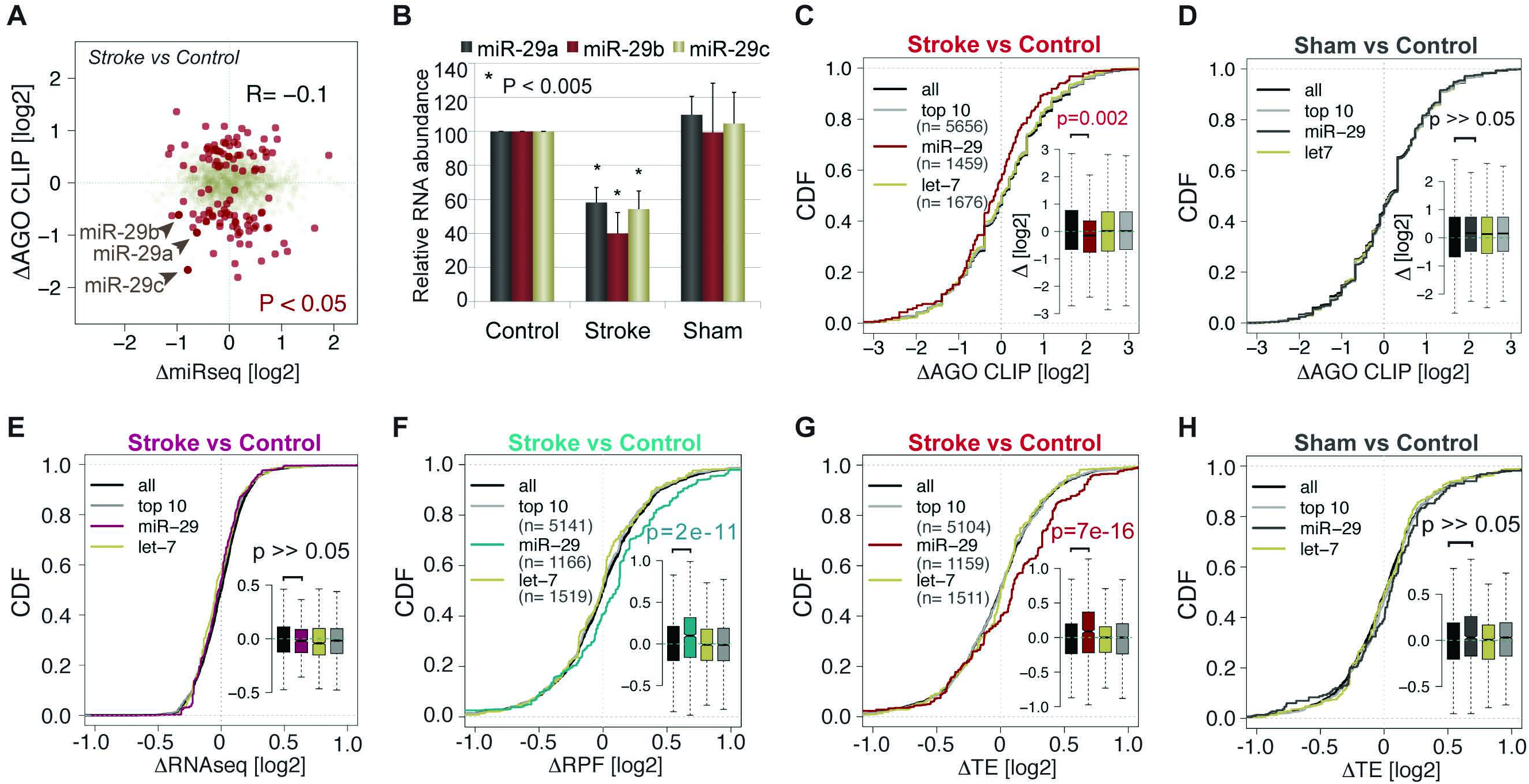
miR-29 is globally de-regulated in stroke. A) Scatter plot of fold change miRNA abundance (x-axis: ΔmiRseq) and AGO-bound miRNA (y-axis: ΔCLIP). Dark red dots indicate significantly changed miRNAs for both (p<0.05). Arrows highlight miR-29 family members. R indicates Pearson’s correlation coefficient. (B) Quantitative PCR show abundance of miR-29 RNAs in stoke or sham cortical tissues normalized to control. P-values generated by Student’s t test. Error bars represent ±SD. Cumulative density function (CDF) plots of (C) stroke and (D) sham AGO CLIP, (E) stroke RNAseq (F) stroke RPF (G) stroke and (H) sham translational efficiency, ΔTE. Curves represent collective changes among miR-29, let-7 or top 10 brain miRNA sites. Inset: box plots with mean fold changes (black notches). P-values calculated using Mann-Whitney-Wilcoxon test. All stroke samples collected after 3hr reperfusion following MCAo. See also Figure S2.

Next, we determined if AGO binding to defined miR-29 sites are lost after I/R by focusing the analysis on canonical 3’UTR binding events. We examined stroke altered peaks with a combined reference data consisting of TargetScan predicted sites (Lewis et al., 2005; Grimson et al., 2007) and empirically identified sites by CLEAR-CLIP on relevant tissue-type, mouse adult cortex (Moore et al., 2015). Cumulative distribution function (CDF) plots of CLIP fold changes showed a significant global reduction in AGO binding to miR-29 sites, while collective binding to highly abundant brain miRNA (‘top 10’) sites and to let-7 sites showed similar distribution as background or ‘all’ sites (Figure 2C). CLIP binding changes across sham and control comparison showed similar distribution for all sites, suggesting down-regulation of miR-29 targets is specific to I/R injury (Figure 2D). These data demonstrate a collective loss of regulation at transcriptome-wide miR-29 target sites following acute I/R injury.

### Acute de-repression of miR-29 targets occurs mainly by translational activation

To assess the functional molecular consequence resulting from global de-regulation at miR-29 sites, we measured steady-state mRNA levels by RNAseq at identical acute time-point as CLIP (3hrs after I/R). miRNA mediated gene repression act mainly through deadenylation and mRNA decay (Guo et al., 2010; Eichhorn et al., 2014). Unexpectedly, miR-29 target mRNA levels did not show noticeable changes in stroke conditions by RNAseq (Figure 2E). This was reflective of global RNA level changes as only 1.3% of genes were found significantly altered upon 3h I/R (300 out of 22,340) (Figure S2C), in direct contrast to the vast transcriptome-wide AGO binding changes detected by CLIP (Figure S2D), suggesting post-transcriptional control by miRNA guided AGO dominate over transcriptional and RNA stabilization mechanisms during acute phase after I/R injury.

We next examined potential translational effects corresponding to the loss of miR-29 control, using a modified ribosomal profiling (RPF) approach to quantify protected mRNA fragments from purified monosomes (Figure S2E) (Ingolia et al., 2009). We saw no difference in the amount of monosomes, genomic distribution of RPF sequencing reads, or read coverage across start and stop codons in stroke condition (3hrs I/R) (Figure S2F-I). However, differential assessment of transcriptome-wide ribosomal association revealed greater global changes in ribosome association in contrast to RNA abundance (Figure S2J-K). Additionally, selective analysis of miR-29 targets showed a robust increase in ribosome association upon I/R injury to control (Figure 2F). Calculating translational efficiency (TE), ratio of RPF density to mRNA abundance (Ingolia et al., 2009), showed even greater increase (Figure 2G). This observation was specific to stroke and not influenced by general surgery procedures, as sham analysis revealed insignificant differences (Figure 2H, S2L). The divergent and orthogonal transcriptome-wide analysis reveal that post-transcriptional regulation by AGO and subsequent translational outcomes dominate to drive physiological response in the acute environment after I/R injury.

### RNA levels of miR-29 targets increase at later stages following stroke

The cellular response to I/R injury is a complicated and dynamic process, involving many factors that generate influence at various times after reperfusion. We asked whether down-regulation of miR-29 and the concomitant activation of its targets are restricted to the initial acute phase following injury. For this, we examined the cortical transcriptome at additional reperfusion time-points (6, 12, and 24hrs after I/R) to generate a time-course miRNA and mRNA profiles (Figure 3A). Time-dependent miRNAseq revealed that miR-29 loss is persistent through 24hrs following I/R. This was true for all three miR-29 family members, while let-7a showed no consistent changes (Figure 3B). Time-course mRNAseq illustrated activation of canonical factors (ie. heat shock proteins and inflammatory cytokine) across all time-points with increasing overall change at later times after I/R (Figure S3A-B). Moreover, in direct contrast to profile at 3hrs, cumulative increases of miR-29 targets were readily detectable at later time-points and correlated with increasing time after I/R (Figure 3C-E). This temporally dependent molecular profile demonstrate the dynamic mechanism of regulatory consequences after I/R injury: the acute and persistent down-regulation of miR-29 initiates a distinct cascade of target activation, exclusively by translational up-regulation at first followed by a steady increase in mRNA levels.

**Figure 3.**
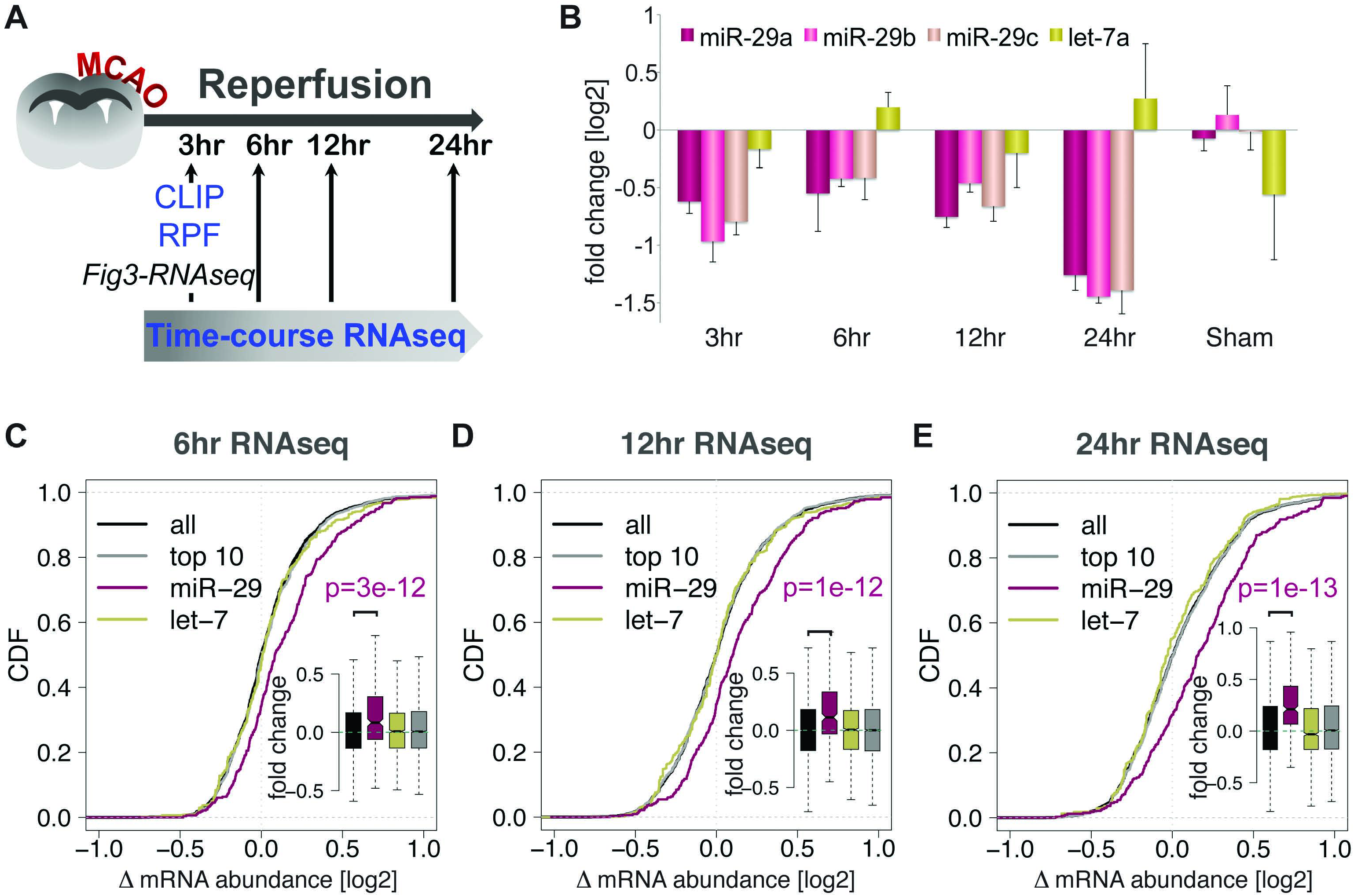
Stroke time-course RNA profiling. (A) Schematic of time dependent transcriptome-wide study of MCAo brain (B) stroke miRNAseq of miR-29 and let-7a at indicated I/R time-points. Error bars represent ±SD. CDF plots of time-course RNAseq for miR-29, let-7, or top10 brain targets at (C) 6hr, (D) 12hr, (E) 24hrs after I/R. Inset: box plots with mean fold changes. P-values generated using Mann-Whitney-Wilcoxon test. See also Figure S3.

### miR-29 target recognition relies on canonical and non-canonical sites

To determine whether miR-29 loss directly activate miR-29 targets, we employed an established *in vitro* system for cerebral ischemic injury. Cultured mouse cortical cells (consisting of ~70% cortical neurons, ~30% astrocytes) were exposed to oxygen-glucose deprivation (OGD). After OGD, the cultures were replenished with full growth media and re-oxygenated to simulate reperfusion. This culture system allows for direct quantification of gene expression and is accessible for loss/gain of function studies (Tasca et al., 2014). The OGD system readily recapitulated the loss of miR-29 measured by qPCR of independent family members (Figure 4A), as well as global activation of miR-29 targets profiled by RNAseq (Figure 4B).

**Figure 4.**
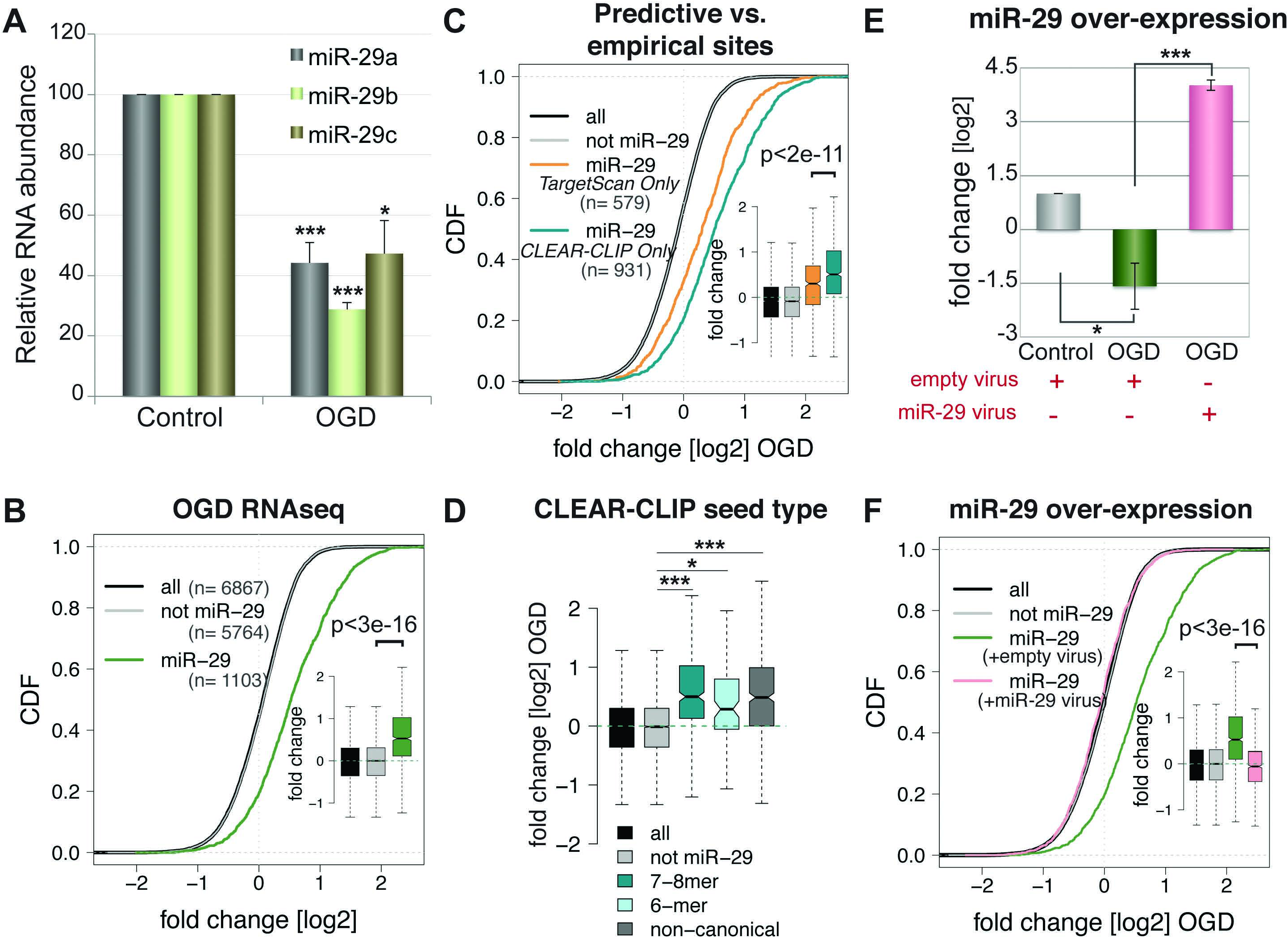
Oxygen-glucose deprivation (OGD) model recapitulates miR-29 de-regulation. (A) Quantitative PCR for miR-29 RNA in OGD treated cortical cultures relative to untreated control, *P<0.005, ***P<5e-6. CDFs of OGD RNAseq for (B) miR-29, and non-miR-29 (‘not miR-29’) targets, (C) miR-29 targets predicted exclusively by TargetScan, or exclusively defined by CLEAR-CLIP. Inset: box plots with mean fold changes. (D) Box plot of OGD RNAseq delineated by miR-29 site seed types. Black notches indicate means. *P<1.7e-11, ***P<2.2e-16. (E) Quantitative PCR show relative fold changes of miR-29 RNA in OGD with or without lentivirus expressed miR-29, normalized to control with empty virus. *P<0.05, ***P<0.005. (F) CDF plot of OGD RNAseq for miR-29 targets infected with empty or miR-29 virus. P-values for all (CDF) calculated using Mann-Whitney-Wilcoxon test, otherwise Student’s t test was used. All error bars represent ±SD.

Unbiased empirical search for *de novo* AGO regulatory events has shown miRNA target recognition can include interactions beyond canonical seed matched pairing (Grimson et al., 2007, Lal et al., 2009, Loeb et al., 2012, Helwak et al., 2013, Moore et al., 2015).

Therefore we compared the function consequences of miR-29 targets defined by TargetScan predicted canonical seed matched sites (7-8mer pairing, highly conserved) and to sites defined exclusively by AGO CLEAR-CLIP (canonical and non-canonical seed matched sites disregarded by computational prediction). CLEAR-CLIP defined miR-29 targets showed greater activation in response to OGD than targets predicted by TargetScan alone (Figure 4C). Further delineation of CLEAR-CLIP defined miR-29 seed types revealed that the greatest response was among targets with 7-8mer seeds and, unexpectedly, among targets containing non-canonical seed sites (Figure 4D), while 6-mer seed targets showed moderate response. These observations suggest a critical role for non-canonical miRNA targets in response to I/R injury and underscore the necessity for both computational and empirical methods to define RNA regulatory networks within relevant physiological systems.

In order to address the requirement for miR-29 loss in target activation, we relied on a gain-of-function experiment using lentiviral gene delivery to over-express miR-29. Quantitative PCR of lentivirus infected cortical cultures showed effective restoration of OGD dependent miR-29 down-regulation (Figure 4E). Accordingly, RNAseq profiles revealed that this restoration was sufficient to suppress global miR-29 target activation (Figure 4F). The loss of miR-29 is a necessary event for the global activation of its targets, as restoration of miR-29 abolishes its transcriptome-wide impact.

### Activated miR-29 targets are enriched in glial genes and activate a regulatory network governing glutamate signaling

In the brain, miR-29 is predominantly expressed by astrocytes, with moderate levels detected in neurons (Smirnova et al., 2005), suggesting a major role for miR-29 in regulating astrocyte physiology. Therefore, we examined miR-29 targets whose expression is also restricted to astrocytes. We ranked our down-regulated AGO peaks by their relative enrichment in astrocytes, using a published reference for brain cell-type transcriptomes (Zhang et al., 2014) (Figure 5A). The top astrocytic miR-29 targets included aquaporin 4 (AQP4; gene symbol: *Aqp4),* known to play a key role in water balance, edema, blood brain barrier permeability and subsequent neuroinflammation in stroke (Fukuda and Badaut, 2012), VEGF, (vascular endothelial growth factor; gene symbol: *Vegfa)* an early hypoxic responder regulating cerebral edema similar to AQP4, switching to promote neurogenesis and vascular repair during later phases (Greenberg and Jin, 2013), and glial mediators of glutamate signaling: mGluR3 (metabotropic glutamate receptor type 3; gene symbol: *Grm3*) and GLT-1 (glial high affinity glutamate transporter, gene symbol: *Slc1a2).* The identification of key astrocytic genes suggests miR-29 targets’ critical brain homeostatic role in responding to I/R injury.

**Figure 5.**
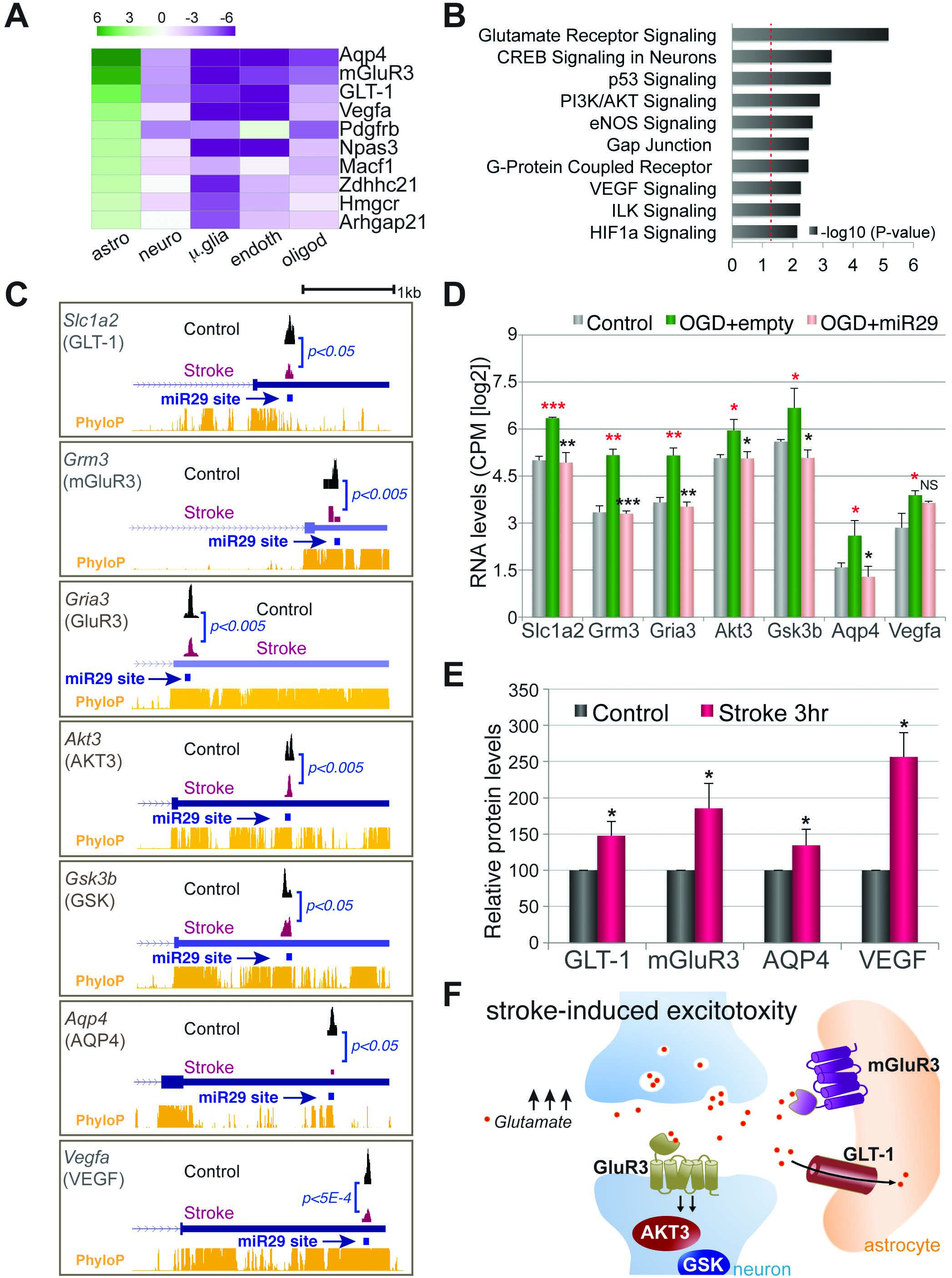
miR-29 targets involved in glutamate signaling. (A) Heat map of top 10 miR-29 guided, down-regulated CLIP targets, ranked by astrocyte enrichment score (B) Pathway analysis, dotted red line indicates p=0.05. C) Genome Browser views for control (black) and stroke (magenta) CLIP peaks. 3’UTRs of 5 representative glutamate genes *(Scl1a2, Grm3, Gria3, Akt3, Gsk3b)* and 2 known stroke responders *(Aqp4 and Vegfa)* shown, miR29 regulatory site indicated by arrows. Yellow track represents conservation (PhyloP) scores. (D) Normalized RNAseq (log_2_ CPM) in control, OGD+empty virus and OGD+miR-29 virus infected conditions. Red Asterisks denote p-values for control vs OGD+empty (red), OGD+empty vs OGD+miR-29 (black). (E) Quantitative westerns for 4 out of 7 representative glial factors of control or stroke cortical tissues following 3hr I/R. (F) Stroke induced glutamate signaling engaging in representative miR-29 targets. *P<0.05, **P<0.005, ***P<5E-4, NS stands for ‘not significant,’ Student’s t test. All error bars represent ±SD. All experiments performed in at least 3 biological replicates. See also Figure S4.

We also performed pathway enrichment analysis for CLIP defined, de-regulated miR-29 targets. Interestingly, the top pathways reflected functions of miR-29 glial targets and included fundamental stress response pathways to oxygen and energy deficit, such as VEGF and HIF1α (hypoxia inducible factor) signaling, and downstream intracellular effector signaling, CREB and PI3K/AKT. Well-known stress signaling pathways, eNOS and p53, were also identified. The most significantly enriched pathway was glutamate receptor signaling (Figure 5B). Strikingly, independent pathway analysis of stroke RPF and OGD RNAseq defined activated miR-29 targets also distinguished glutamate signaling (S4A-B), providing robust and orthogonal support for enrichment of glutamate response genes among miR-29 targets. Glutamate excitotoxicity is a primary cause of tissue damage elicited by I/R stress. A sudden ischemic attack and corresponding depletion of cellular ATP causes a sudden K^+^ efflux and induce depolarization. This ‘anoxic depolarization’ accumulates excitatory signals (ie. glutamate) to overwhelm synaptic load and cause neuronal death (Moskowitz et al., 2010). Acute activation of targets by a de-regulated miRNA may prompt a protective response to control glutamate signaling in an effort to overcome excitoxicity.

We further examined individual targets involved in glutamate signaling and validated their response to the loss of miR-29 regulation. The representative glutamate genes included glial enriched glutamate transporter, GLT-1, glial specific glutamate receptor, mGluR3, neuronal glutamate receptor, GluR3 (ionotropic AMPA3 receptor; gene symbol: *Gria3),* and downstream effectors of activated signaling AKT3 (gene symbol: *Akt3*) and GSK (gene symbol: *Gsk3b).* We also included *Aqp4* and *Vegfa*, miR-29 targets and established stroke response genes, in our assessment. Genome visualization of stroke AGO CLIP data showed significant decrease in respective 3’UTR peaks at corresponding miR-29 sites for all representative genes (Figure 5C). Quantitative expression analysis of demonstrated RNA accumulation in OGD and most importantly, miR-29 over-expression repressed induced RNA levels to near baseline for 6 out of 7 genes (Figure 5D). Moreover, quantitative assessment of selected glial factors, GLT-1, mGluR3, AQP4 and VEGF, confirmed their increase in steady-state protein levels within cortical stroke tissues early (3hr) upon I/R injury (Figure 5E, S4C). Taken together, the data reveal an acutely activated miRNA target network in response to anoxic depolarization and excess glutamate. Responsive miR-29 targets respond to glutamate signals by engaging post-synaptic and glial-specific glutamate receptors (GluR3 and mGluR3), in addition to downstream signaling effectors (AKT3 and GSK). Remarkably, a critical glial factor for uptake of excess glutamate (GLT-1) is also involved, indicating an effort to counteract excitoxicity and to restore brain glutamate homeostasis (Figure 5F).

### miR-29 regulated reuptake signal controls brain glutamate levels

GLT-1 (excitatory amino acid transporter 2, EAAT2) is a major glutamate transporter in the brain, playing critical roles in buffering glutamate levels. We further investigated the role of miR-29:GLT-1 regulatory axis in mediating glutamate homeostatic functions in I/R injury. Interestingly, the AGO down-regulated miR-29 site on GLT-1 3’UTR does not contain canonical seed matched sequence. We therefore tested the functionality of this site by design of a target site blocker (TSB) to CLIP defined 20nt AGO binding region (Figure 5C, *Slc1a2*). As expected, miR-29 over-expression in OGD cultures (+control TSB) repressed OGD dependent accumulation of GLT-1 RNA, however, TSB against miR-29 binding site of GLT-1 3’UTR abrogated miR-29 repression to near baseline levels, as GLT-1 RNA levels were similarly induced as in OGD without miRNA overexpression (Figure 6A). This suggests that precise 3’UTR recognition by miR-29 is required for GLT-1 RNA regulation in OGD. To determine if miR-29:GLT-1 interaction has an effect on GLT-1’s ability to uptake extracellular glutamate, we quantified glutamate concentrations in OGD culture media in presence or absence of miR-29 and TSB (Figure 6B). OGD stimulus readily increased glutamate levels in media, and overexpression of miR-29 by lentiviral gene delivery exacerbated this induction. Intriguingly, site-specific inhibition of miR-29:GLT-1 by TSB counteracted the miR-29 dependent glutamate accumulation (Figure 6C). The data suggests miR-29 de-regulation and responsive activation of GLT-1 may play a role in cellular glutamate regulation by promoting uptake of toxic glutamate in response to I/R injury.

**Figure 6.**
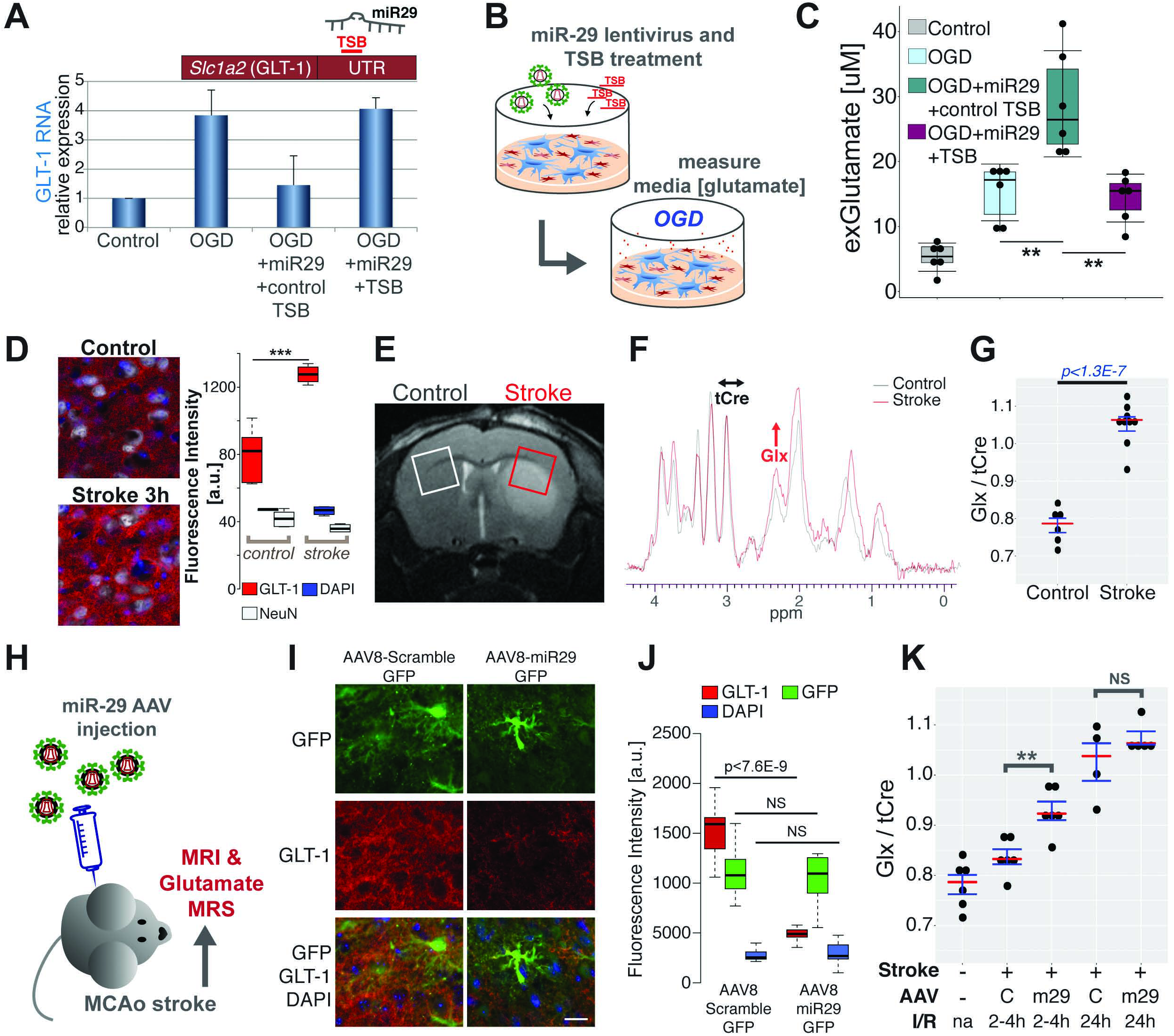
miR-29:GLT-1 regulatory axis control glutamate levels. (A) Quantitative PCR of GLT-1 RNA in OGD or control, with miR-29 lentivirus, and/or TSB (B) Schema of media glutamate assay to test miR-29:GLT-1 regulatory function in OGD. (C) Extracellular glutamate quantification. (D) Immunofluorescent staining and quantification of local GLT-1 protein in ischemic periphery (MCAo cortical tissue) for control and 3hr after stroke. (E) T_2_ – weighted image of MCAo brain. Boxes specify voxels placed for ^1^H-MRS data acquisition for each hemisphere (F) ^1^H-MRS spectra of stroke (red) and control (grey), arrows indicate corresponding Glx (red) or tCre (black) peaks. (G, K) Quantification of relative local glutamate (Glx/tCre) (H) Schema of MRS experiment using AAV-miR-29 injected MCAo animals. (I) GLT-1 staining of cells infected with AAV8-GFP scramble control or miR-29. Scale bar = 20μm. (J) Quantification GFP, GLT-1, and DAPI fluorescent signals. *P<0.05, **P<0.005, ***P<5E-4, NS stands for ‘not significant,’ Student’s t test. All error bars represent ±SD. All experiments performed in at least 3 biological replicates. See also Figure S5.

We took this investigation further, to demonstrate *in vivo* evidence of miR-29 control on GLT-1 and glutamate in stroke animals. Local GLT-1 protein levels in cortical tissue surrounding ischemic territory are robustly up-regulated upon stroke (3hrs post I/R), suggesting localized activation of glutamate re-uptake signal (Figure 6D). Proton magnetic resonance spectroscopy (^1^H-MRS) is a non-invasive clinical approach to monitor and quantify levels of brain metabolites altered in disease (Ramadan et al., 2013). We leveraged this technology to assess brain glutamate levels in stroke MCAo animals. ^1^H-MRS spectra reliably measured brain glutamate associated Glx peak, resonating between 2.2-2.6ppm (Figure S5A). Data acquisition for each ^1^H-MRS scan requires a precise placement of a 2mm cubic voxel at a localized region of interest. Hence, we relied on T2 MRI acquired images of ischemic territory to appropriately focus our measurements near I/R damaged regions (Figure 6E). Using this method, we successfully detected a significant induction of brain glutamate in stroke hemisphere compared to control (Figure 6F-G). For testing miR-29 regulated glutamate in I/R injured brain, we engineered adeno-associated virus (AAV) to deliver miR-29 by stereotaxic injections. MCAo surgery was then performed on injected animals for assessment by MRI/MRS (Figure 6H). GFP reporter encoding AAV was successfully delivered within desired brain region (Figure S5B). In addition, by using an AAV serotype characterized for preferential infection of astrocytes, AAV8 (Aschauer et al., 2013), we were able to effectively target gene delivery to glia, allowing a robust signal to noise assessment of astrocyte-specific GLT-1 (Figure S5C). We also confirmed miR-29 regulation of GLT-1 in cortical stroke tissue, as AAV8-miR-29 robustly silenced GLT-1, while scramble control virus had no effect (Figure 6I-J). Remarkably, ^1^H-MRS measurements revealed a significant increase in brain glutamate levels animals injected with AAV-miR29 over scrambled controls (Figure 6K). This observation was exclusive to early times (2-4hr) upon I/R. At a later time (24hr) we detected extremely high levels of glutamate in both control and in treated animals and saw no effects with AAV-miR29 pre-treatment. Thus activated GLT-1 is sufficient for attenuating glutamate levels early upon injury, however, not a later time when excitotoxic effects reach overwhelming levels. This temporal dynamic of stroke brain glutamate levels reflects our global observation of miR-29 target activation (main Figures 2–3). Acute post-transcriptional effects dominate early molecular responses and govern immediate physiological outcomes such as glutamate control. While later consequences mediated by increase in target RNAs may facilitate secondary and chronic responses to I/R insult. Collectively, these results serve as strong *in vitro* and *in vivo* evidence for an activated miRNA target facilitating homeostatic control in I/R injured brain.

### Identification of human genetic polymorphisms in stroke-associated miR-29 sites

To apply our findings from animal model system to human stroke, we generated human brain AGO CLIP data to define orthologous, stroke-associated, human miRNA regulatory sites. For human CLIP, we used post-mortem medial prefrontal cortex tissues from 6 individuals. To further increase biological complexity for robust peak calling, we merged our in-house generated data with a human AGO CLIP resource (Boudreau et al., 2014) generated from motor cortex and cingulate gyrus of 17 individuals. Together, these data defined 23,662 reproducible (peak height >50) human AGO brain peaks. Intersecting these sites to mouse stroke peaks (999 Hg19 mapped out of 1,190) identified 429 orthologous AGO regulatory sites (Figure 7A). Peak evaluation and motif analysis revealed human brain enriched miRNAs (Figure 7B), including miR-29 and another stroke associated miRNA, miR-181 (Ouyang et al., 2012).

**Figure 7.**
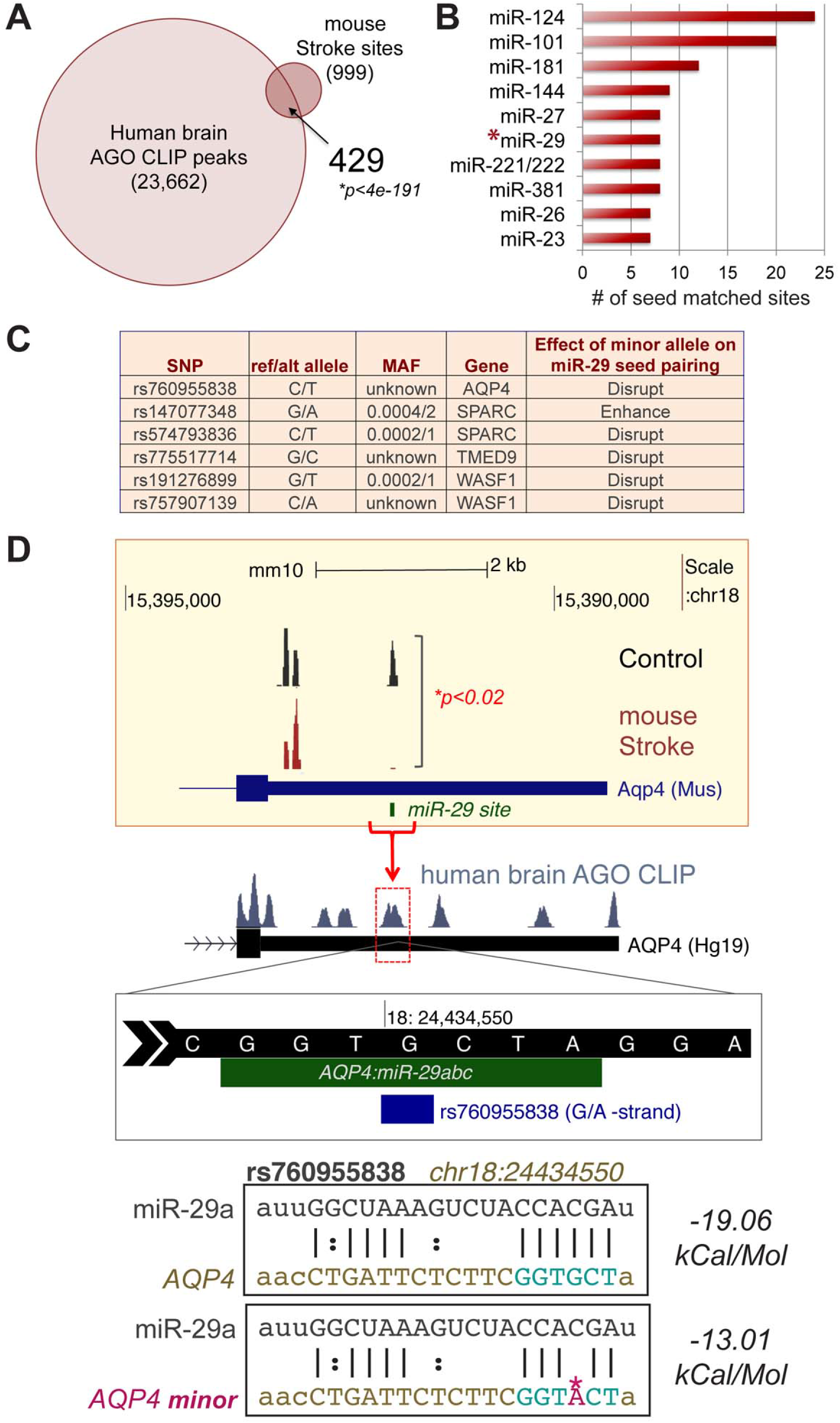
Stroke-associated miR-29 target site polymorphisms. (A) Venn diagram of intersected stroke associated human miRNA regulatory sites. *Hypergeometric test was used to calculate p-value. (B) Top 10 miRNA seeds enriched among 429 regulatory sites. Asterisk highlights miR-29. (C) Table of SNPs in stroke-associated miR-29 sites, reference or alternative nucleotide allele, minor allele frequency (MAF), gene name, and predicted influence on seed pairing. (D) Genome browser views of AQP4 3’UTR, mm10 (top), Hg19 (below). AGO CLIP stroke peak is bracketed. Red arrow/dotted square indicate orthologous miR-29 site in human AQP4. rs760955838 is positioned within miR-29 seed recognition sequence. Predicted energy coefficients for miR-29a pairing (bottom panel). See also Figure S6.

The human genome contains thousands of variants that can potentially alter miRNA binding. Allele specific polymorphisms in miRNA binding sites have been shown to vary miRNA function and play essential roles in complex human diseases, including cancer, diabetes, and Alzheimer’s diseases (Brewster et al., 2012, Zhao et al., 2013, Delay et al., 2011). We implemented an informatic analysis to identify single nucleotide polymorphisms within the 429 stoke-associated miRNA sites. We relied on existing SNPs previously submitted to NCBI’s dbSNP resource and intersected them with 7-8mer seed containing, miRNA binding sites within our 429 cohort. We identified 332 unique single nucleotide variations among 165 sites representing 222 miRNAs across 117 genes (Supplementary Table 5). Notably, we identified 6 SNPs that modify miR-29 target sequences. The alternative alleles among these 6 sites represent both rare and common variants within the human population. 5 out of the 6 variants have the potential to disrupt miR-29 binding, while 1 may enhance miR-29 seed pairing (Figure 7C, S6). Some reside within a single 3’UTR polymorphisms for two distinct miR-29 binding sites (rs147077348 and rs574793836, SPARC 3’UTR), and others alter divergent nucleotides within a single miR-29 seed binding sequence (rs191276899 and rs757907139, *WASF1* 3’UTR). Interestingly, we identified a SNP within a known stroke response gene (rs760955838, AQP4 3’UTR). rs760955838 is a rare variant, the alternative allele (A) indicating a lower predicted binding energy coefficient than reference allele (G), thus may weaken miR-29 seed pairing and affect *AQP4* gene expression (Figure 7D). This study illustrates a unique approach to investigate disease relevant polymorphisms altering RNA regulatory functions and may provide useful genetic resource for better understanding clinical stroke outcomes.

## DISCUSSION

Empirical investigation of unique biological and pathological states is imperative for elucidating mechanisms underlying relevant physiology. AGO CLIP affords the power to systemically identify and quantify miRNA:target interactions in normal and disease brain, prevailing informatic predictions. In this study, AGO-CLIP allowed us to discover acutely activated network of miRNA targets due to a de-regulated miR-29 in I/R injury. The dynamic molecular mode of global activation is led by a strong translational effect with minimal impact on mRNA levels. Such temporally precise regulatory action of miRNA, limited to distinct physiological circumstances have been similarly described in the observation that translational effects precede mRNA turnover in exerting miR-430 control during early phases of zebrafish development (Bazzini et al., 2012). Additionally, reminiscent of molecular responses seen underlying axonal regeneration and repair upon neuronal injury (Jung et al., 2012), the immediate translational activation of miR-29 targets maybe the most pertinent response to modulate the acute and rapid cellular responses required for counteracting ischemic insult.

Unexpectedly, AGO-CLIP, interpreted together with brain CLEAR-CLIP data, revealed that non-canonical seed pairing sites contributed to many functional miR-29 regulatory events. Such finding reinforced the power of empirically derived datasets unafforded by those exclusively reliant on predictive strategies (ie. TargetScan) generated from irrelevant tissue types and relies on evolutionary constraints for interpretation. Auxiliary target interactions at miRNA’s 3’ ends give distinction among miRNA family members with shared seeds and also can compensate imperfect seed pairing for miRNAs with AU-rich seed motifs (Moore et al., 2015, Broughton et al., 2016). Although miR-29 seed sequence is not AU-rich, further characterization of compensatory pairings among identified seedless interactions may yield mechanistic insights distinct from canonical miRNA targeting and help delineate gene ‘sub-networks’ playing disparate roles in I/R injury.

We uncovered a novel regulatory network of miR-29 targets facilitating key astrocytic functions to regain homeostatic control upon I/R injury, including maintenance of blood brain barrier and glutamate signaling. Astrocytes play a central role in brain homeostasis, particularly via cooperative processes they establish with neurons to supply energy metabolites, mediate neurotransmitter action, and regulate water and ion concentrations (Phatnani and Maniatis, 2015). We find astrocytic miR-29 targets, along with few neuronal targets, engage glutamate signaling in a consorted effort to buffer toxic glutamate levels, as altering miR-29 levels effect brain glutamate accumulation. We find that this is in part due to miR-29 specific regulation of GLT-1 (EAAT2), a glial-specific glutamate transporter that control glutamate levels by its rapid uptake from extracellular space. Moreover, the precise miR-29 regulatory site on GLT-1 3’UTR determines glutamate concentration. Interestingly, miR-29 effect on brain glutamate is exclusive to acute time-points in I/R injury, reflecting time-sensitive global mechanism of target activation and may suggest distinct physiology mediated by targets activated at later time-points.

Previous studies of miR-29 in stroke animal models show improved outcomes upon altering miRNA levels (Khanna et al., Wang et al., 2015). However insightful, these studies were limited by examinations of single miRNA:target interactions, thus only providing an isolated perspective on global miRNA function. Furthermore, physiological effects of activated targets are not singular, as some facilitate neuroprotective functions while others exhibit deleterious ones. This complexity underscores the importance of examining network effects of transcriptome-wide regulation at distinct I/R time-points to gain full perspective of the dynamic, systems-level response to I/R injury.

Stroke-relevant miR-29 sites identified by AGO CLIP also provided a powerful source for determining genomic variants with potential associations to disease. Interestingly, miR-29 de-regulation has been observed in more archetypal neurodegenerative diseases, such as in Alzheimer’s and Huntington’s effected brains, and in sciatic nerve injury response (Hébert et al., 2008, Johnson et al., 2008, Verrier, et al., 2009), suggesting an expansive and crucial role for miR-29 in brain homeostasis. A comprehensive, unbiased characterization of miR-29 targets in various disease states will further provide insights into both common and distinct pathological gene networks, help elucidate universal miR-29 target site variants associated with risk or outcome for multiple neurodegenerative diseases, and allow elegant designs for therapeutics of broad benefit.

## ACKNOWLEDGEMENTS

We thank Darnell and Iadecola lab members for invaluable suggestions, expertise, and feedback, Henning Voss and Eric Aronowitz of Weill Cornell Medicine Citigroup Biomedical Imaging Center for MRI and MRS help, Rockefeller Genomics Resource Center and New York Genome Center for hi-throughput sequencing. Support provided by the National Institutes of Health (NS034389, NS081706, NS097404, and 1UM1HG008901 to R.B.D, NS34179 to C.I., NS081179 to J.A., and NS067078 to P.Z.). Additional support was provided by the Simons Foundation (SFARI 240432) to R.B.D, Renal Research Institute to J.P.D., and Feil Family Foundation to C.I. R.B.D. is an Investigator of the Howard Hughes Medical Institute.

## AUTHOR CONTRIBUTIONS

M.K., J.A., C.I., and R.B.D conceived the project. M.K. designed all studies and interpreted results. C.B. and C.P. performed MCAo surgeries. C.A. performed OGD experiments. K.U. performed stereotaxic injections. M.J.M provided mouse brain CLEAR-CLIP data. J.P.D. performed MRS data analysis. M.K. and A.M. generated AGO CLIP datasets. M.K. and J.J.F generated miRNAseq and RNAseq datasets. C.Y.P. ran miRANDA algorithm for SNPs. M.K. performed all other experiments, informatic analysis, and made figures. M.K. and R.B.D wrote the manuscript with input from all authors. R.B.D, C.I., J.A., and P.Z. supervised the research, and provided resources.

---

**Figure S1.**
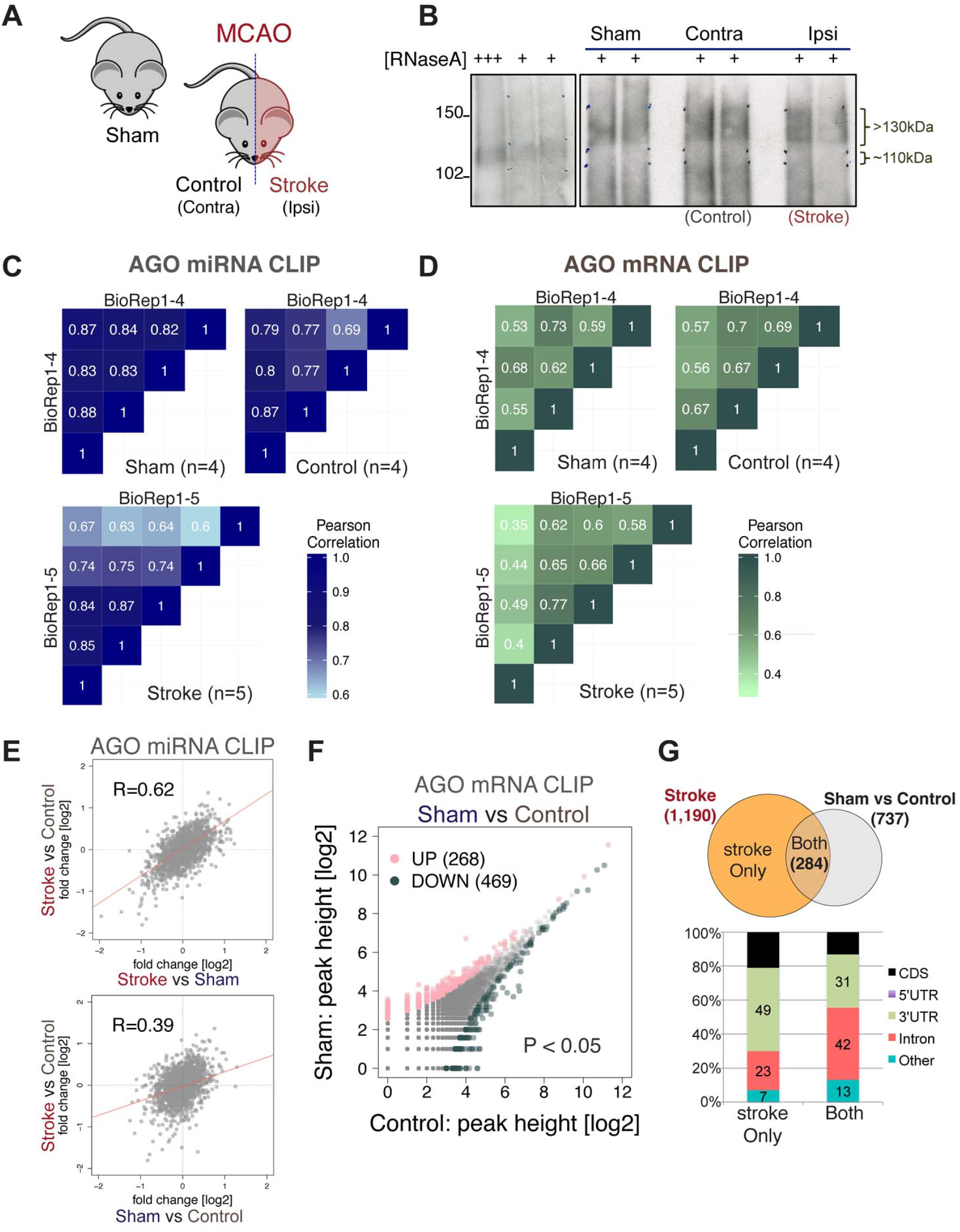
Kobayashi et al.

**Figure S2.**
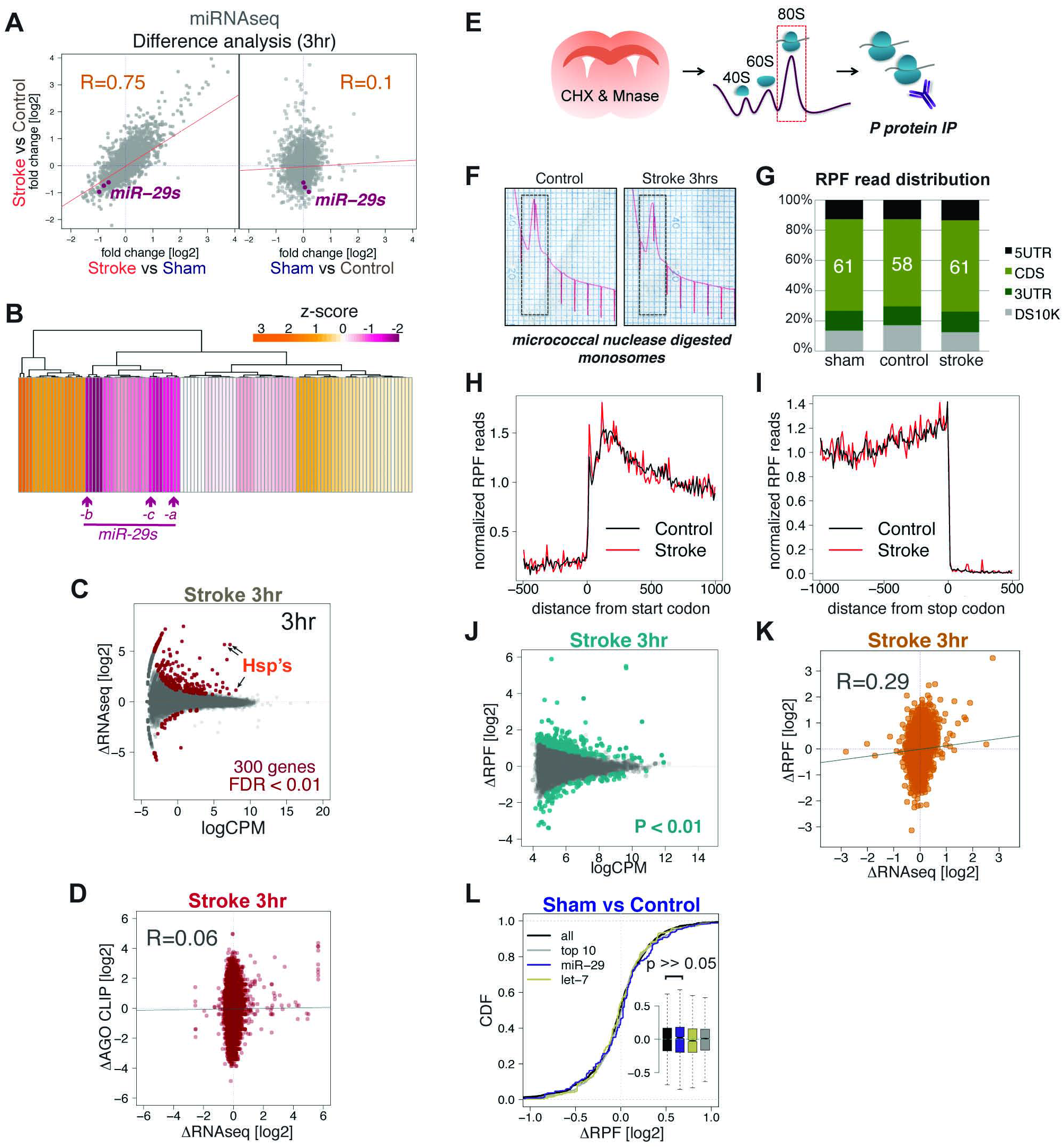
Kobayashi et al.

**Figure S3.**
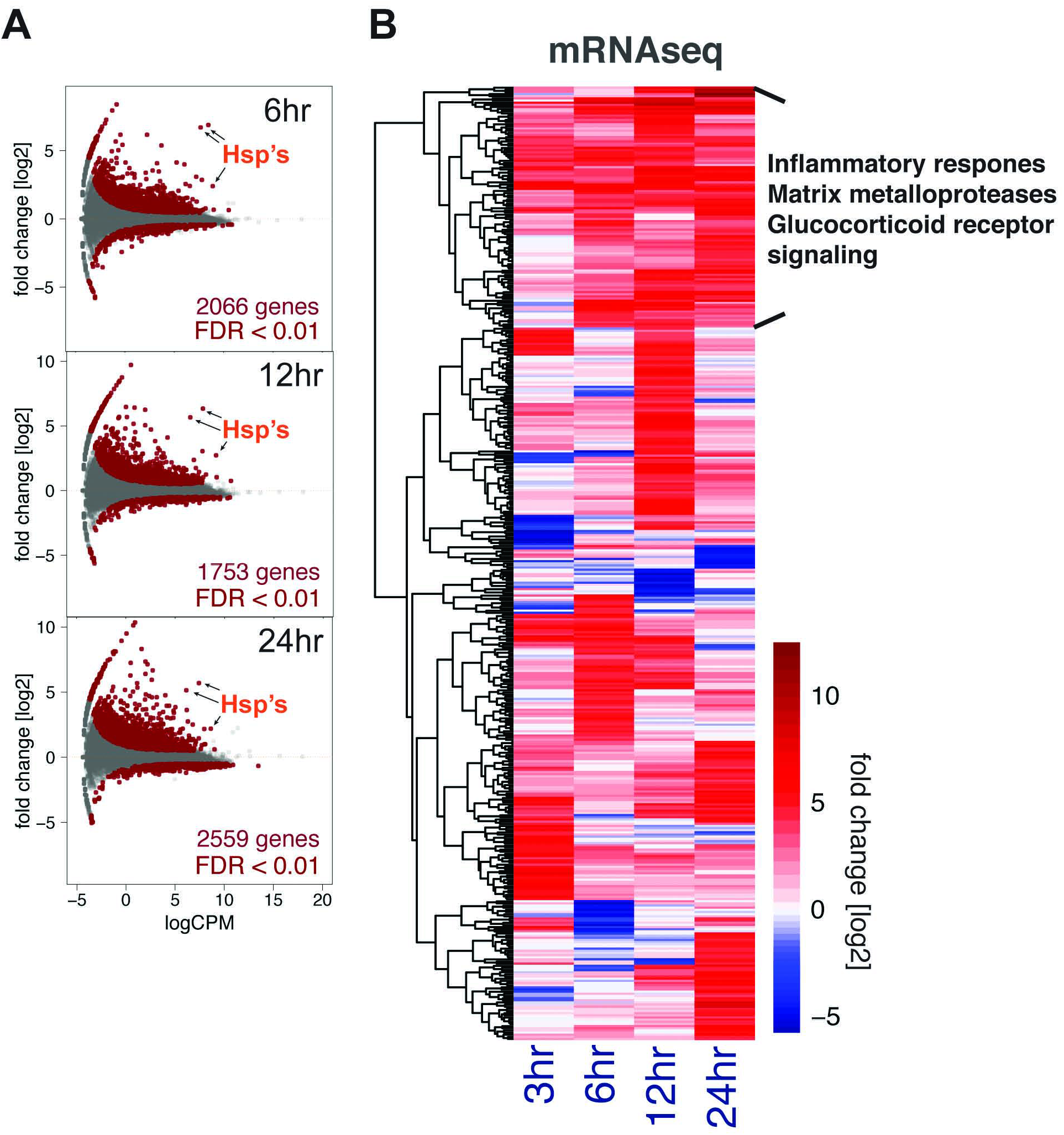
Kobayashi et al.

**Figure S4.**
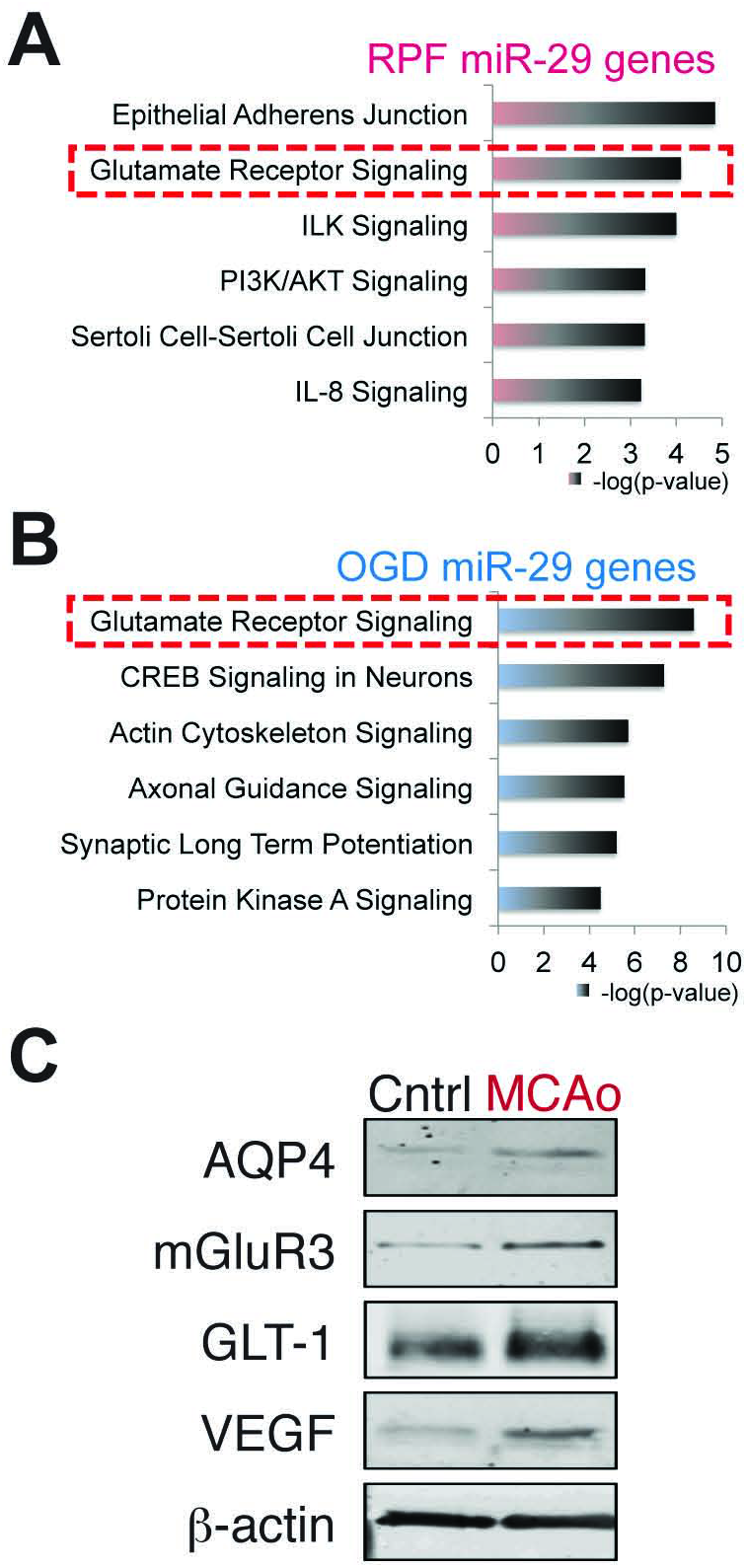
Kobayashi et al.

**Figure S5.**
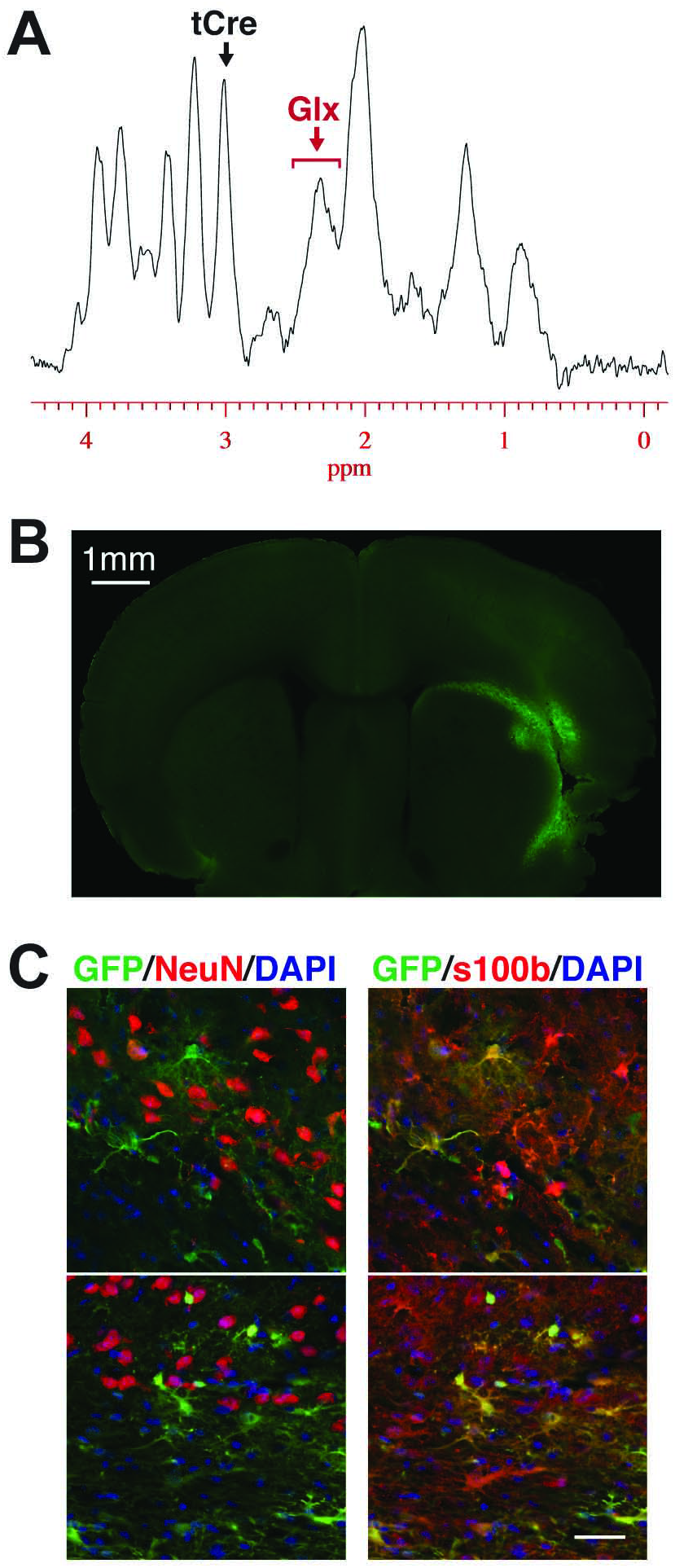
Kobayashi et al.

**Figure S6.**
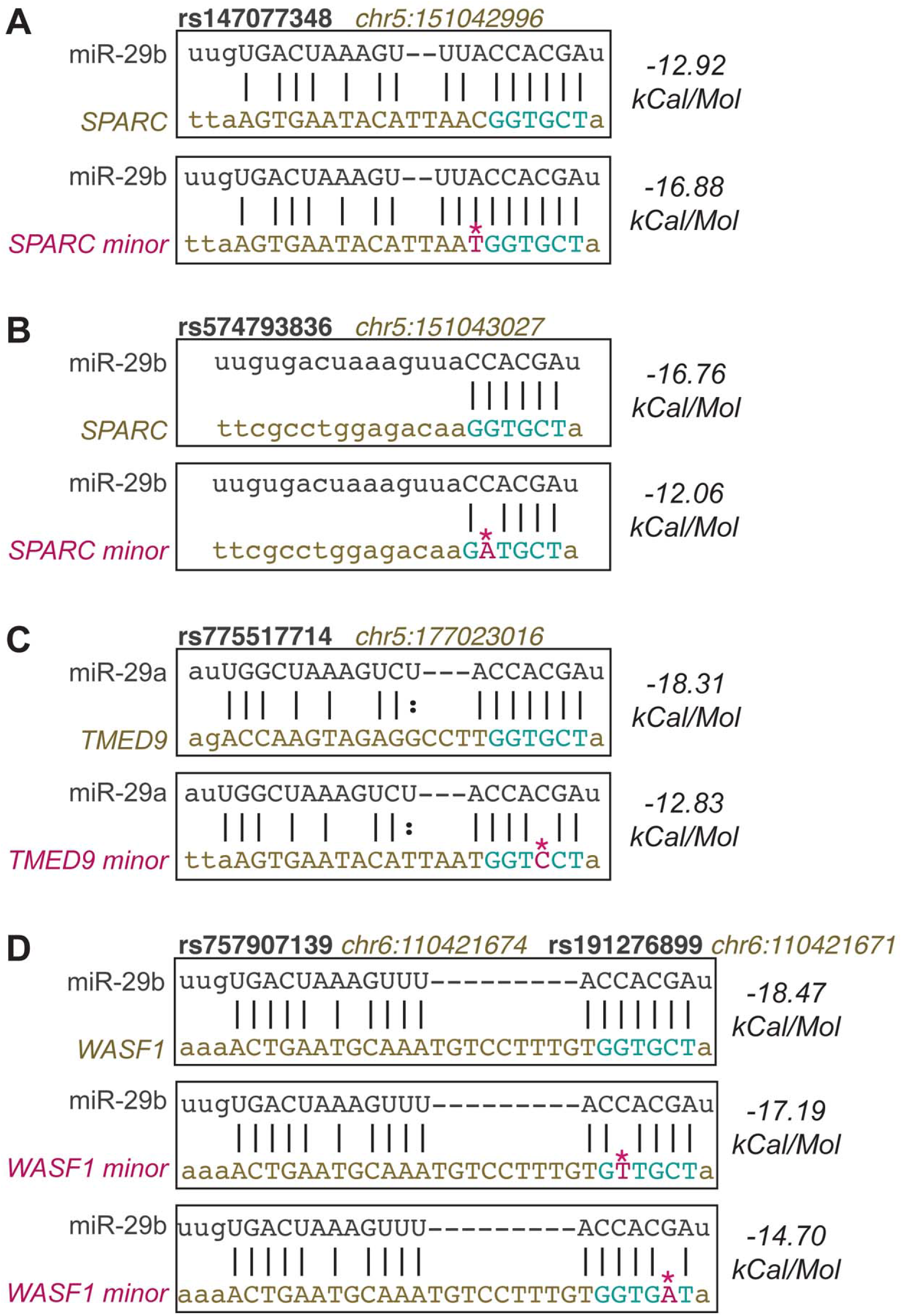
Kobayashi et al.

